# Analysis of a detoxified *Escherichia coli* strain for bacteriophage production

**DOI:** 10.64898/2026.04.21.719556

**Authors:** Emily A. Welham, Alba Park de la Torriente, Jia Arng Lee, Marianne Keith, Sean McAteer, Gavin K. Paterson, David L. Gally, Alison S. Low

## Abstract

Phage therapeutics are re-emerging as adjuncts or alternatives to antibiotics and their clinical translation will be enhanced with production methods that minimise downstream processing. We evaluated whether an endotoxin-reduced *E. coli* strain developed for production of recombinant proteins, ClearColi®, can serve as a useful, safe phage production host without compromising yield and whether targeted receptor complementation can expand its utility. The parent strain BL21(DE3), and its lipid A modified derivative, ClearColi®, were compared with respect to infection and generation of phage. Across a panel of 31 phage, a similar host range was observed between BL21(DE3) and ClearColi®. To expand host range *ompC* was genetically engineered into the chromosome of ClearColi®, thereby adding OmpC-dependent phage to its production capacity. Production metrics were broadly comparable between the hosts; efficiency of plating and final titres for representative phage were not significantly different; burst size varied by phage but without consistent host bias. Endotoxin activity in ClearColi®-propagated lysates was reduced by over 1000-fold relative to BL21(DE3), reaching the low hundreds of endotoxin units (EU) versus hundreds of thousands for BL21(DE3). Intravesical administration of ClearColi®-derived phage (LUC4) into pigs elicited no clinical abnormalities and no significant increases in circulating cytokines up to 48 hours after administration. ClearColi® allows efficient production of diverse phage with low endotoxin, reducing the requirement for downstream processing. Although its minimal LPS reduces its capacity for producing some LPS-dependent phage and its growth is slower than BL21(DE3), requiring optimisation for maximal phage titre, the safety and simplified manufacturing process support further development of endotoxin modified strains for phage production.

**Impact statement:** Antibiotic resistance is a current global problem and treatments based on phage and phage products already have a proven track record with particular bacterial infections, especially in the urinary tract. While progress is being made on *in vitro* phage synthesis, large scale bacteriophage preparations require a bacterial host for production, consequently toxic components in the initial lysate need to be removed or significantly diluted for safe clinical use. This is a study of the potential to utilise an endotoxin-reduced *E. coli* strain, ClearColi®, to produce safer phage therapeutics. Such endotoxin modified strains should minimise the processing steps required and reduce overall production costs of a phage preparation. The research demonstrates that the endotoxin-reduced strain was able to produce a wide range of phage and for studied examples at phage titres equivalent to the more toxic parent strain. We also show that the strain can be modified to increase its host range and confirm the very low endotoxicity of basic phage lysates produced by the strain. Replicating this process to engineer additional low-toxicity bacterial production strains will accelerate the development of safer, more cost-effective phage therapeutics.

## Introduction

Bacteriophage (phage) are abundant viruses that exploit the cellular machinery of their bacterial hosts to replicate and produce progeny (1). The rise in multidrug resistance (MDR) has reignited interest in phage therapies. Whether used as an alternative or an adjunct treatment to traditional antimicrobials, these viruses have shown therapeutic potential in treating infections (2).

Currently phage production is more commonplace in agriculture and the food industry than in clinical settings. Multiple products are available to reduce bacterial burden across diverse sectors, from aquaculture to fresh produce and dairy (3–5). Historically, countries in Eastern Europe have been developing phage for clinical use since their discovery by Twort (1915) and d’Hérelle (1917), whilst Fleming’s discovery of penicillin led to the overshadowing of phage therapies elsewhere (6,7). However the current era of antimicrobial resistance (8) has led to a resurgence in the West of interest in developing phage therapeutics. Regulatory hurdles still prevent the use of phage as a mainstream treatment and its use is often limited to extenuating circumstances, as a compassionate last attempt when other treatments have failed. The reported efficacy of phage treatments varies widely reflecting divergent success rates among personalised therapeutics, general applications and the combinatorial strategies with antibiotics (9).

Despite their potential, the development of phage-based therapies is hindered by the inherent complexity of regulating biological entities and difficulties in ensuring consistent safety, scalability and batch-to-batch reproducibility. Phage generally require a host bacterial strain to be produced at the concentrations required for treatment, and this means any bacterial debris that may be an issue for human or animal health, have to be removed from the primary phage lysate. In the case of the Gram-negative bacteria, such as *Escherichia coli* (*E. coli*), this includes removal of the highly inflammatory endotoxin, a component of the lipopolysaccharide (LPS) that forms part of the outer membrane (10). In the UK, the Medicines and Healthcare products Regulatory Agency (MHRA) have developed a series of regulatory conditions for production of phage therapeutic products (11). Developing well-characterised, low-endotoxin productions strains would fundamentally enhance the safety and traceability of Gram-negative phage products.

LPS is comprised of three regions; the O-antigen (polysaccharide), the core unit and lipid A/endotoxin (12). Lipid A, is the amphipathic moiety which anchors lipopolysaccharide in the bacterial cell’s outer membrane (13). Endotoxin contains a varying number of fatty acid chains which trigger conformational change in mammalian produced toll-like receptor 4 complexed to myeloid differentiation factor 2 (TLR4-MD2), in turn triggering intracellular signalling pathways (14,15). It is the predominant inflammatory factor of Gram-negative bacteria and can trigger an excessive inflammatory response, or cytokine storm, leading to vascular permeability and septic shock in humans (16,17). Phage therapeutics produced from Gram negative bacteria undergo endotoxin removal processes such as ultracentrifugation, Triton X phase separation and anion-exchange chromatography and post-production quality control methodologies are required to quantify remaining endotoxin (10,18,19). As per current guidance, a maximum concentration of endotoxin which can be safely administered intravenously is 5 EU kilogram^-1^ hour^-1^ (20,21).

An *E. coli* phage production strain with a detoxified lipid A moiety would allow the production of therapeutics that require less post-production purification. A derivative of the laboratory strain BL21(DE3) (DE3), named ClearColi®, has reduced endotoxin activity and is commercialised for the production of recombinant proteins (19). The ClearColi® strain was developed from a *kdsD* and *gutQ* deleted strain (KPM22 L11), which contained a *msbA* suppressor mutation allowing growth (19). Unmarked deletions of genes *lpxL, lpxM, pagP lpxP and eptA* ensure that only lipid*IV*_A_, precursor can be produced, to keep some membrane stability whilst removing the majority of endotoxicity (19). It is noted that even small amounts of endotoxin left in a sample can trigger human primary CDlc^+^ dendritic cells and as such a NF-kB reporter assay, using cells overexpressing TLR4 is recommended to identify endotoxin activation of TLR4, or the industrially standardized limulus amoebocyte assay (LAL) to identify endotoxin presence (22).

ClearColi® served as a proof of principle for low-endotoxin phage production, yielding phage lysates with comparable performance to a representative BL21(DE3) parent lineage strain. By genetically engineering a phage receptor into the ClearColi® genome, we successfully expanded its host range. Crucially, intravesical instillation of these lysates in a porcine model elicited no systemic innate immune response, demonstrating the safety and clinical viability of this production platform.

## Methods

### Strain Information

*E. coli* KPM404, engineered to only express endotoxin precursor Lipid IV_A_, was developed from BL21(DE3) and obtained as ClearColi® by RTC and Lucigen Corp. ClearColi® contains deletions of *kdsD, gutQ, lpxL, lpxM, pagP, lpxP* and *eptA* along with a suppressor mutation in *msbA148* (19). It is used for academic research purposes only. BL21(DE3) from our laboratory stocks, was originally obtained from Novagen. *E. coli* O25b:H4-ST131, strain EC958 (23) was used from laboratory stock originally provided by Prof Matthew Upton, University of Plymouth. Master stocks were stored at −70°C in 25% glycerol.

### Growth Media and Culture Conditions

Lysogeny (Luria) Broth, Lennox (LB) (Sigma-Aldrich) and LB-Agar (Miller, Formedium) were used for culturing bacterial stocks. In general, 5mL LB was inoculated with a single *E. coli* colony and incubated at 37°C overnight, shaking at 170 rpm. Overnight cultures were sub-cultured by diluting 1:100 into prewarmed LB and cultured in the same conditions to the required optical density (OD_600_) for use in experiments.

### Phage isolation and propagation

Phage used were provided by the Phage Technology Centre, GmbH, Bonen, Nordrhein-Westfalen, Germany or isolated in our laboratory from wastewater samples provided by the Scottish Environment Protection Agency. Phage were propagated in an appropriate host strain in LB, prepared and titred as previously described (24). Sterile Saline-Magnesium (SM) Buffer (100 mM NaCl, 8 mM MgSO4, 50 mM Tris–HCl (1M, pH 7.5), and 0.01% (w/v) gelatine was used to store phage at 4°C.

### Bacterial Growth Rates and Doubling Times

Fresh cultures of BL21(DE3) and ClearColi® were cultured as above. Overnight cultures were sub-cultured by diluting 1:100 into prewarmed LB in flasks and incubated at 37°C, shaking at 180 rpm, for 6 hours. OD_600_ was measured at 30-minute intervals.

Growth rates were calculated using; 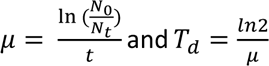 where µ = growth rate and T_d_ = doubling time. N_0_ = initial OD_600_ reading, N_t_ = final OD_600_ reading, t = time of exponential phase between N_0_ and N_t_.

### Interaction Assays and Area Under the Curve (AUC) Analysis

Phage interaction assays were performed with *E. coli* isolates ClearColi® and BL21(DE3) using 31 phage at an MOI of 0.01 (1:100, phage: bacteria) as previously described (24). Phage interaction assays were completed as three biological repeats each with three technical replicates, with growth recorded over 18 hours and data analysed using GraphPad PRISM (11.0.0) and area under the curve (AUC) scores calculated.

### Bacteriophage Receptor Definition

Using natural mutagenesis to select naturally generated single nucleotide polymorphisms (SNPs) in the bacterial genome, phage receptors were defined in permissive host strains. 100 µL of the permissive strain and 100 µL neat (n>1×10^8^ PFU/mL) phage was added to 3mL 0.7% LB Top Agar and layered onto an LB agar plate. Incubation at 37°C overnight allowed phage resistant colonies to develop, these were picked and streaked out three times for purification. A phage rechallenge validated resistance and the bacterial mutant was sent to MicrobesNG (Birmingham, UK) for Illumina sequencing. Illumina reads were aligned with wildtype GenBank file of the parent strain to determine which SNPs, or small-scale deletion/insertions were selected. Snippy (25) was run on the Galaxy.eu server to identify which mutations conferred phage resistance, with the assumption that in most cases these encode or regulate expression of the essential phage receptor.

### Efficiency of Plating

Comparative efficiency of phage plating on strains; BL21(DE3) and ClearColi® were compared. LB agar plates with 3mL 0.7% Top Agar seeded with respective bacterial strains had 10 µL of serially diluted phage diluent plated. Plates were incubated at 37°C overnight before plaque forming units were counted, biological triplicates were completed with the 31 phage. PFU/mL vs *E. coli* strain were equated and plotted, in GraphPad Prism, to compare plaquing efficiencies.

### Burst Size Experiments

The burst size from BL21(DE3) and ClearColi® with phage (HAM53, B01, BAT1 and N) were compared. Overnight bacterial cultures were prepared as previously described and plaques counted on BL21(DE3). BL21(DE3) and ClearColi® were diluted 1:100 into 10 mL LB with 2mM CaCl_2_, incubated at 37°C x180 rpm until a working OD_600_ =0.5 was reached. 9.9mL bacterial cells and 100 µL phage (1×10^7^ PFU/ml) were added in Erlenmayer flasks and incubated together for 5 mins in a 37°C water bath. An absorbance control was taken from one flask, with 1mL taken into a 1.5mL Eppendorf containing 50µL chloroform (Serva Electrophoresis GmbH) and vortexed. 1mL was transferred to another flask, and 100µL serially transferred to a second and then third flask. Flasks were sampled at regular timepoints by adding 100µL flask culture to 100 µL BL21(DE3) plating host and mixed into 3 mL top agar, layered onto LB agar plates. Samples were taken for a minimum of 1 hour. Plates were incubated overnight at 37°C before counting of PFU and analysis of data in GraphPad Prism.

### Restoration of ompC expression

BL21(DE3), and other *E. coli* B strains, reportedly lack functional *ompC* (26). Wildtype *ompC* from *E. coli* strain EC958 was restored in the ClearColi® genome using a well-documented allelic exchange method (27). The verification of sensitivity was confirmed, using an *ompC*-dependent phage, LUC4 (24), to determine if ClearColi®+*ompC* could now be infected. Primers used found in **Table 1**.

**Table 1.**
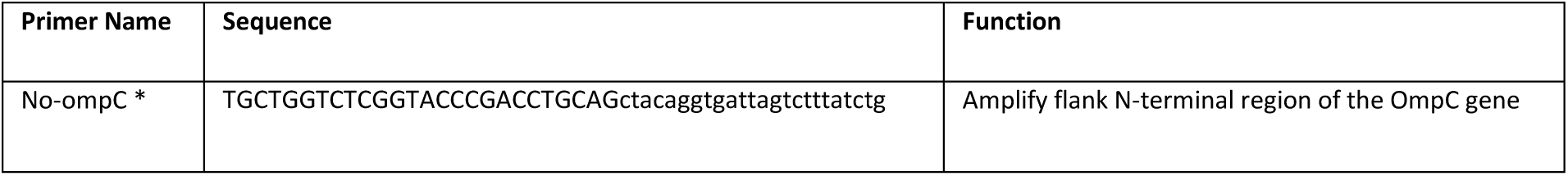
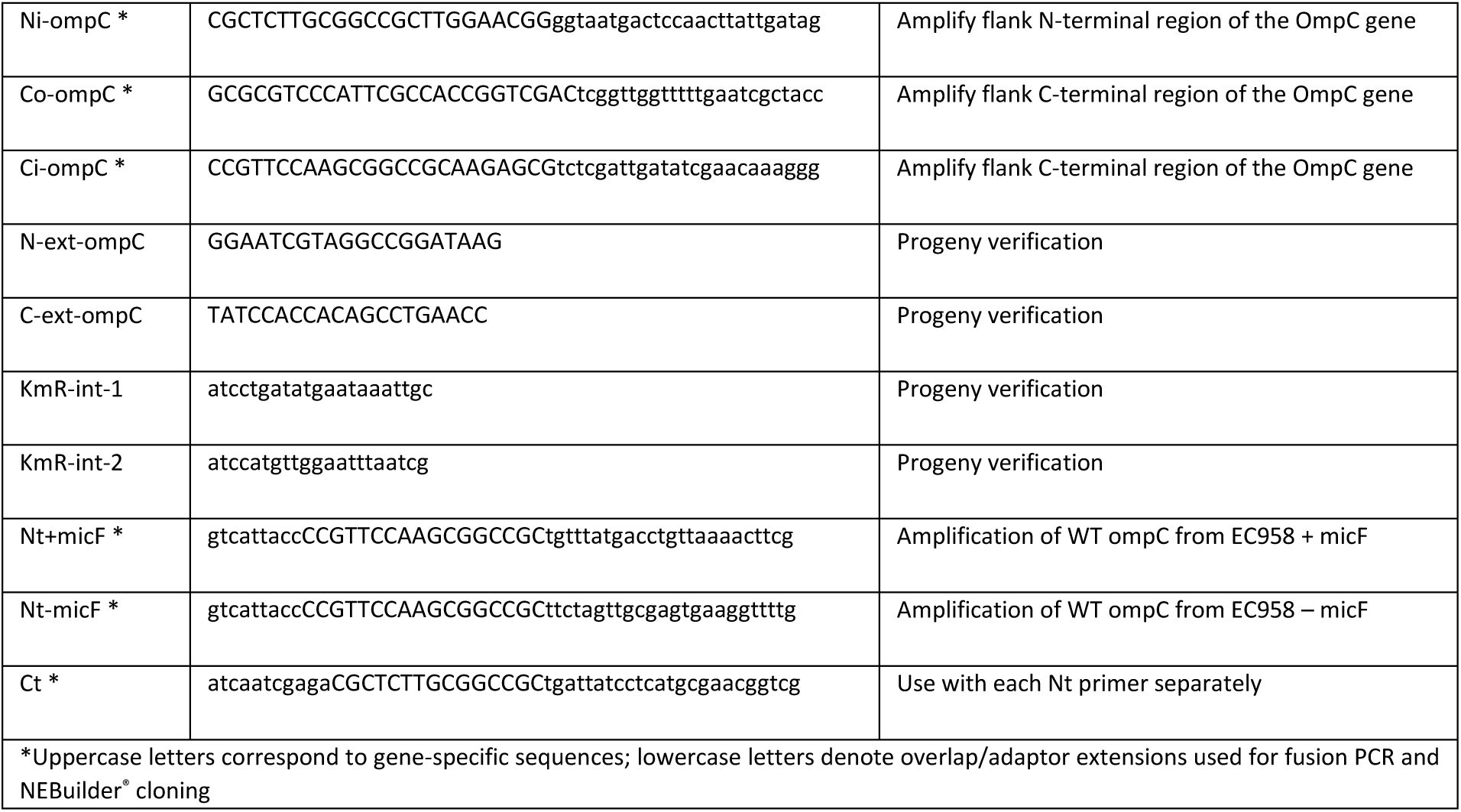
Primer sets used in restoration of *ompC* gene into ClearColi®.

### Endotoxin Detection Assays

HEK-Blue-4 is a cell-based assay using HEK293 cells engineered to express TLR4. Endotoxin triggers the TLR4 response in the mammalian immune response, thus the cells detect endotoxin signalling via the NF-κB pathway, leading to expression of secreted embryonic alkaline phosphatase (SEAP) as a reporter. SEAP develops into a deep blue/purple colour on addition of Quanti-Blue dye. The endotoxin concentration within 0.01-0.1 EU/mL is proportional to absorbance (OD_600_) of the sample. This HEK-Blue LPS Detection kit 2 (InvivoGen, USA) was used as per the manufacturer’s guidance. (28–33)

### LUC04 Production and use in a Porcine Safety Trial

Phage LUC4 was propagated in ClearColi®+*ompC* in order to generate a preparation with low levels of endotoxin. This was tested using the HEK cell-based endotoxin activity assay as above. The LUC4 preparation had an endotoxin content of ∼100 EU (endotoxin units)/ml, 1 EU is equal to 0.1 ng of active *E. coli* endotoxin (34,20,21). Body weights of pigs ranged between 40 - 45kg, so 1-part neat phage was diluted in 50-parts medical grade 0.9% NaCl solution and 100 ml doses were prepared with a final phage titre of 1.37 x 10^9^ PFU/ml and a total of 200 EU units. Stability, infectivity of phage, pH and sterility were measured after 48 hours storage at 4°C. Blood samples were taken prior to phage instillation into the bladder then 100 mL of LUC4 preparation was slowly administered, via transurethral catheter into three juvenile pigs under general anaesthesia. The pigs remained under anaesthesia with the catheter occluded for 30 minutes post procedure to minimise immediate expulsion of the phage preparation. The pigs were monitored closely for two hours post-instillation and then observed at least twice daily. Further blood samples were taken under sedation at 24 hours and at 48 hours just prior to euthanasia.

### Measurement of inflammatory markers from blood samples

The blood samples were centrifuged at 100 RCF for 5 minutes to pellet the red blood cells. The serum was harvested, aliquoted and stored at −80°C until use. Samples were defrosted and cytokine levels measured using a Cytokine & Chemokine 9-Plex Porcine PorcartaPlexTM in a Magpix System (Luminex) following the assay protocol in the user guide. Each sample was run in duplicate.

## Results

### ClearColi® and the BL21(DE3) strain maintained in our laboratory exhibit distinct growth kinetics

While ClearColi® is derived from a BL21(DE3) lineage, our laboratory BL21(DE3) strain represents a separate lineage rather than the direct progenitor of the commercially sourced ClearColi® used in this study. To validate previously reported differences in growth (19) 6-hour growth curves were generated for both strains used in this study (**Fig. 1a**). The laboratory BL21(DE3) strain demonstrated a highly reproducible exponential growth phase with a calculated doubling time of 27.7 ± 2.08 minutes. In contrast, ClearColi® exhibited a shallower exponential gradient resulting in an increased doubling time of 47.7 ± 6.43 minutes (**Fig. 1b**). This growth disparity was also reflected on solid media; after 24 hours at 37 °C, BL21(DE3) formed typical creamy/white round colonies on LB agar, while ClearColi® appeared only as tiny ‘pinprick’ colonies (**Figs 1b and 1c**). Although ClearColi® colonies became clearly visible following an additional 24-hour incubation, they remained substantially smaller than those of the parent lineage, confirming a markedly attenuated growth rate on a rich medium.

**Fig 1.**
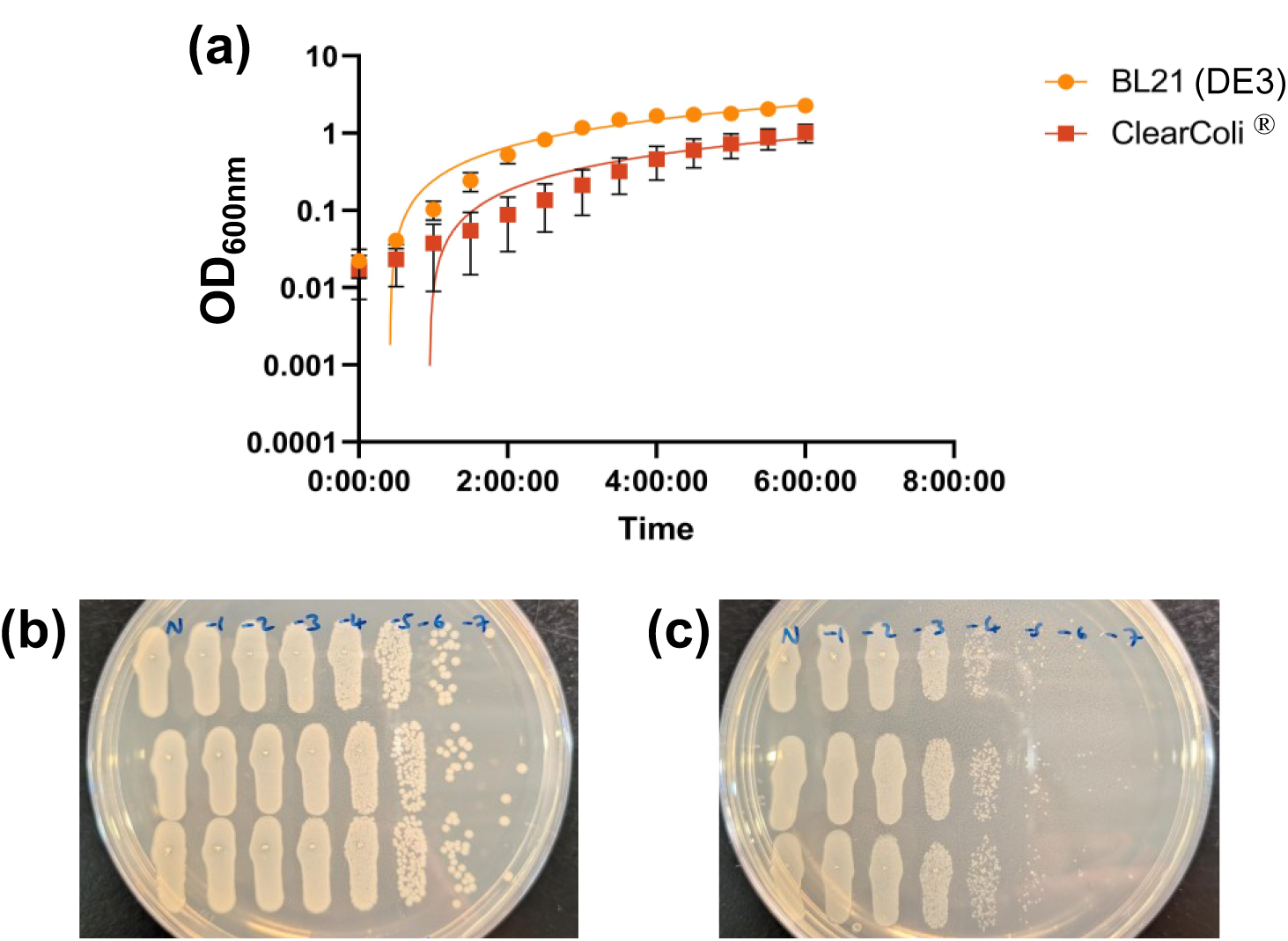
Growth of ClearColi® compared to strain BL21(DE3). **a)** 6-hour bacterial growth curves, with standard curve used to calculate growth rate and subsequent doubling time of BL21(DE3) and ClearColi®. Data points represent the average of three biological replicates. **b)** Ten-fold serial dilution of BL21(DE3) to show colony formation after 24 hours incubation at 37°C. **c)** Ten-fold serial dilution of ClearColi® to show colony formation after 24 hours incubation at 37°C.

### ClearColi® and BL21(DE3) exhibit similar phage susceptibility profiles

In an 18-hour growth assay across a panel of 31 diverse phage, both strains showed similar infective patterns, with phage activity scored using an Area Under the Curve (AUC) calculation and these scores compared between the two host strains (**Fig. 2 and Table S1**). Only 3 phage; LUC4, AB19 and GEO showed less activity in ClearColi® compared to BL21(DE3). A score of 60 or less was defined as an active phage (24). BL21(DE3) was susceptible to 22 phage, while ClearColi® was susceptible to 20. From the panel, 12 phage are predicted to utilise protein receptors, 1 is predicted to use capsule, 14 are predicted to require lipopolysaccharide (LPS) and the rest remain uncharacterised (**Table S1**). Given that BL21(DE3) possesses a truncated LPS lacking the O-antigen (19), and ClearColi® produces only the precursor Lipid*IV*_A_ reduced infection by LPS-targeting phage was anticipated. The two strains shared nearly identical resistance profiles but of the 8 unable to infect either host, 4 were predicted to require LPS and 1 predicted to require capsule, also likely to be missing from these laboratory derived strains (**Table S1**).

**Fig. 2.**
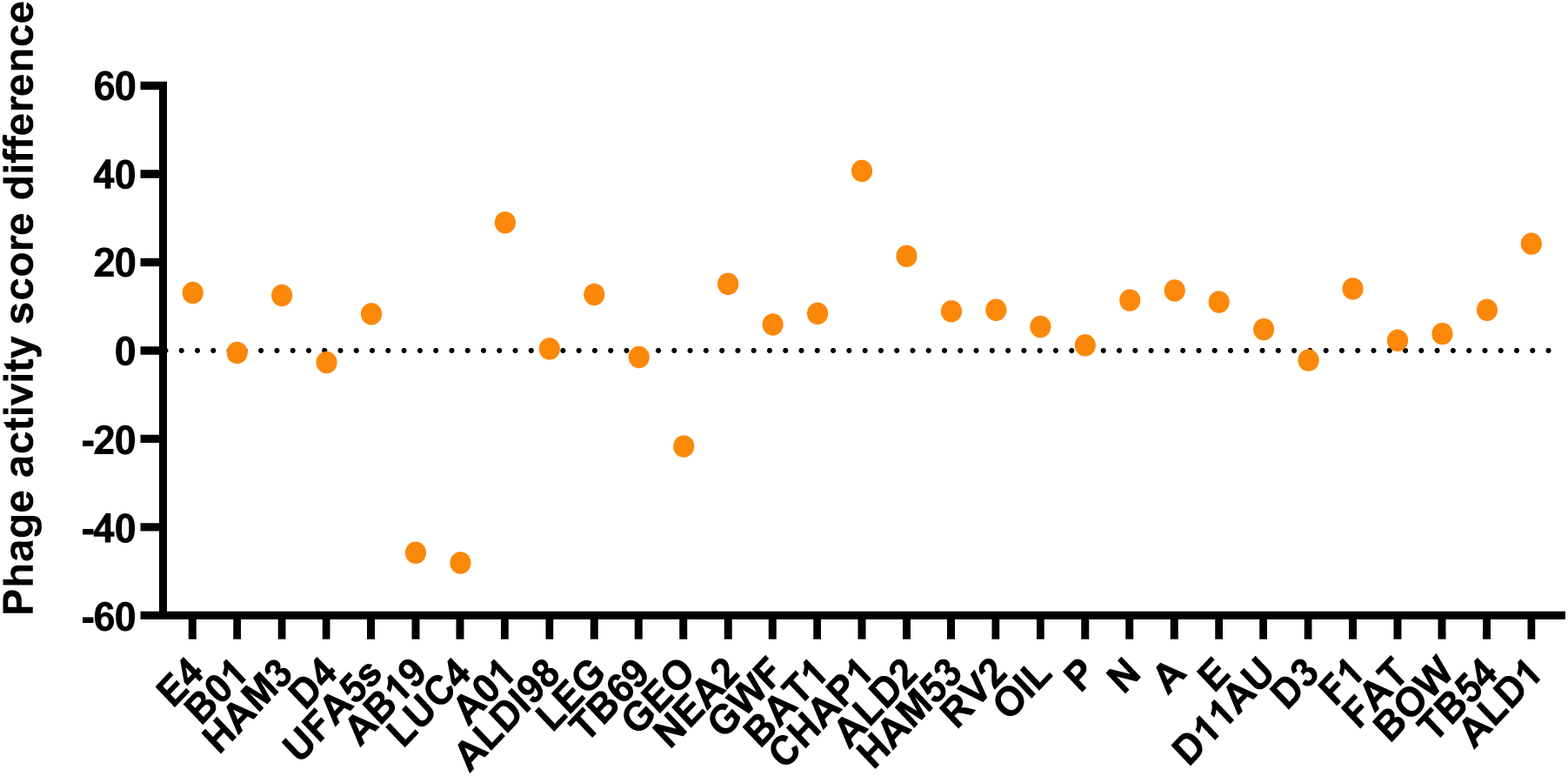
Comparison of phage infectivity between ClearColi® and BL21(DE3). Difference between BL21(DE3) and ClearColi® phage activity scores (using AUC of growth over 18 hours) are shown for each phage (x-axis). Negative values represent less phage activity on ClearColi® compared to BL21(DE3), and positive values show more activity.

### ClearColi® performs similarly to BL21(DE3) as a phage production host

To evaluate the utility of ClearColi as a production host its performance against BL21(DE3) was compared using Efficiency of Plating (EOP), final phage titres and burst size. The EOP was calculated for the phage that had measured infective activity in both strains in liquid media. Across the phage panel the EOP was largely equivalent between the parent lineage and the derivative, though some inter-assay variation was observed across biological replicates (**Fig. 3a**). From this panel, four distinct phage (N, BAT1, B01 and HAM53) were selected for further characterisation. Lysate enumeration on their respective host-seeded top agar revealed no significant differences in titres produced by either strain (**Fig. 3b**). Comparative burst size analysis revealed phage-specific replication dynamics between the two hosts (**Fig. 3c**). While phage BO1 demonstrated near identical burst sizes across both strains, significantly different outcomes were observed for phage N and phage BAT1. Interestingly, phage N exhibited a significantly larger burst size in the ClearColi® background (*P* = 0.04), whereas BAT1 production was better in the parental BL21 (DE3) lineage (*P* = 0.01). For phage HAM53, although the mean burst size appeared higher in ClearColi®, the difference was not statistically significant due to high biological variance. Collectively this data suggests that whilst the burst sizes were inherently variable and phage-dependent, they were not consistently higher in one strain over the other. Notably the differences in burst size did not appear to correlate with final phage concentrations. Collectively these data indicate ClearColi® performs as a robust alternative to standard BL21(DE3) for high-titre phage production.

**Fig. 3.**
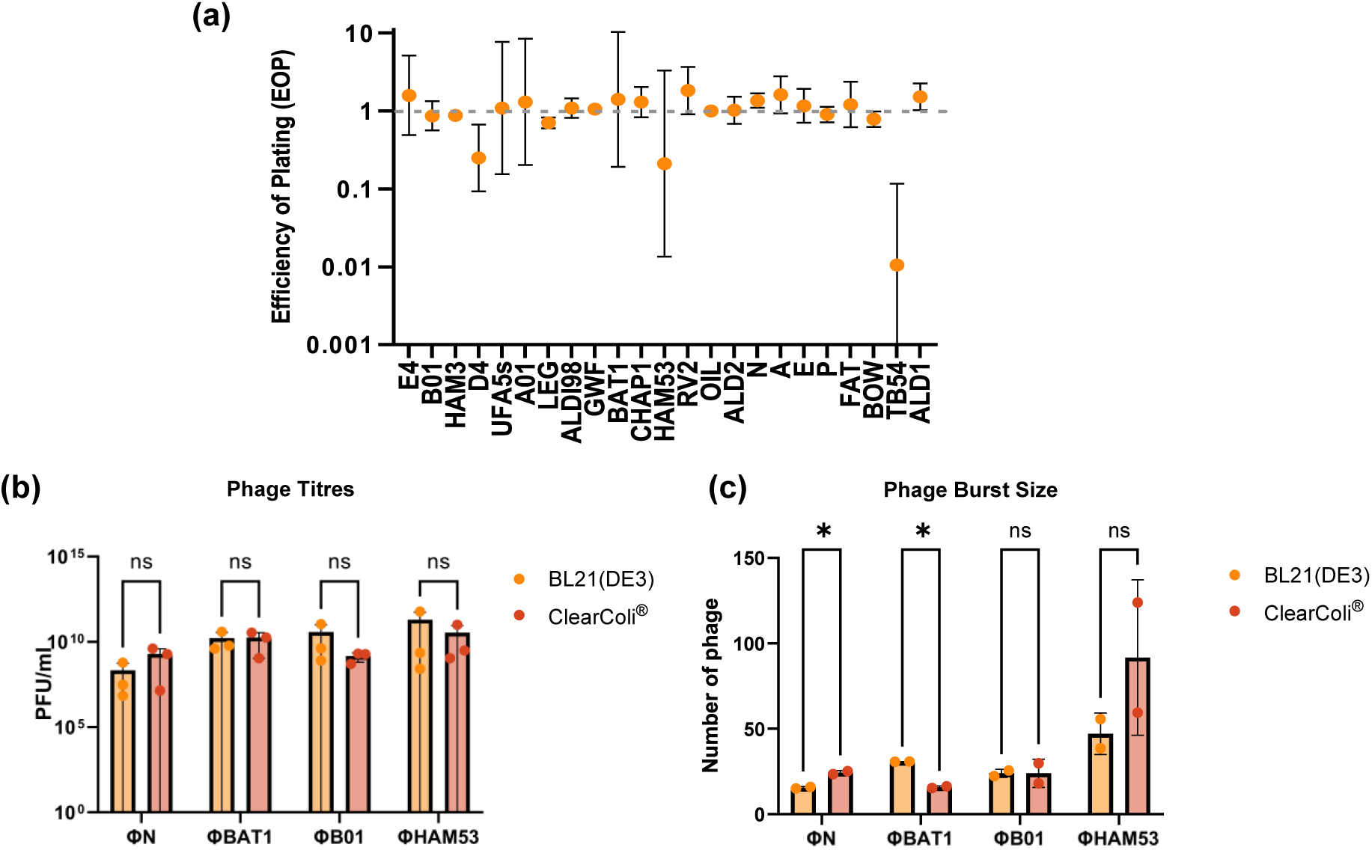
Production capacity of ClearColi® compared to BL21(DE3). **a**) Efficiency of Plating; Scores < 1 represent more plaques forming on BL21(DE3) compared to ClearColi®. Scores = 1 have equivalent plaques form on BL21(DE3) and ClearColi®. Scores > 1 represent more plaques forming on ClearColi®. **b)** Phage titres present no significant difference in production number of virions, using a multiple unpaired t test. **c**) Phage burst sizes by host calculated from two biological replicates of one-step growth curves, mean PFU ± standard deviation plotted. Statistical significance calculated by a multiple unpaired t-test analysis.

### Endotoxin content of phage lysates is significantly reduced when ClearColi® is used as a production host

Original laboratory phage stocks were generated in each phage’s respective wild-type host strain necessitating a two-stage propagation strategy to mitigate endotoxin carryover. While primary ClearColi**®** lysates showed significant endotoxin reductions compared to BL21(DE3) controls by HEK-Blue-4 assay (**Fig. S1**), residual variance suggested minor contamination from original host stocks. To isolate the effect of the production strain a second round of propagation was performed (**Fig. 4**). This resulted in uniform, very low measured endotoxin levels across all four phage, representing a thousand-fold reduction compared to BL21(DE3) derived lysates.

**Fig. 4.**
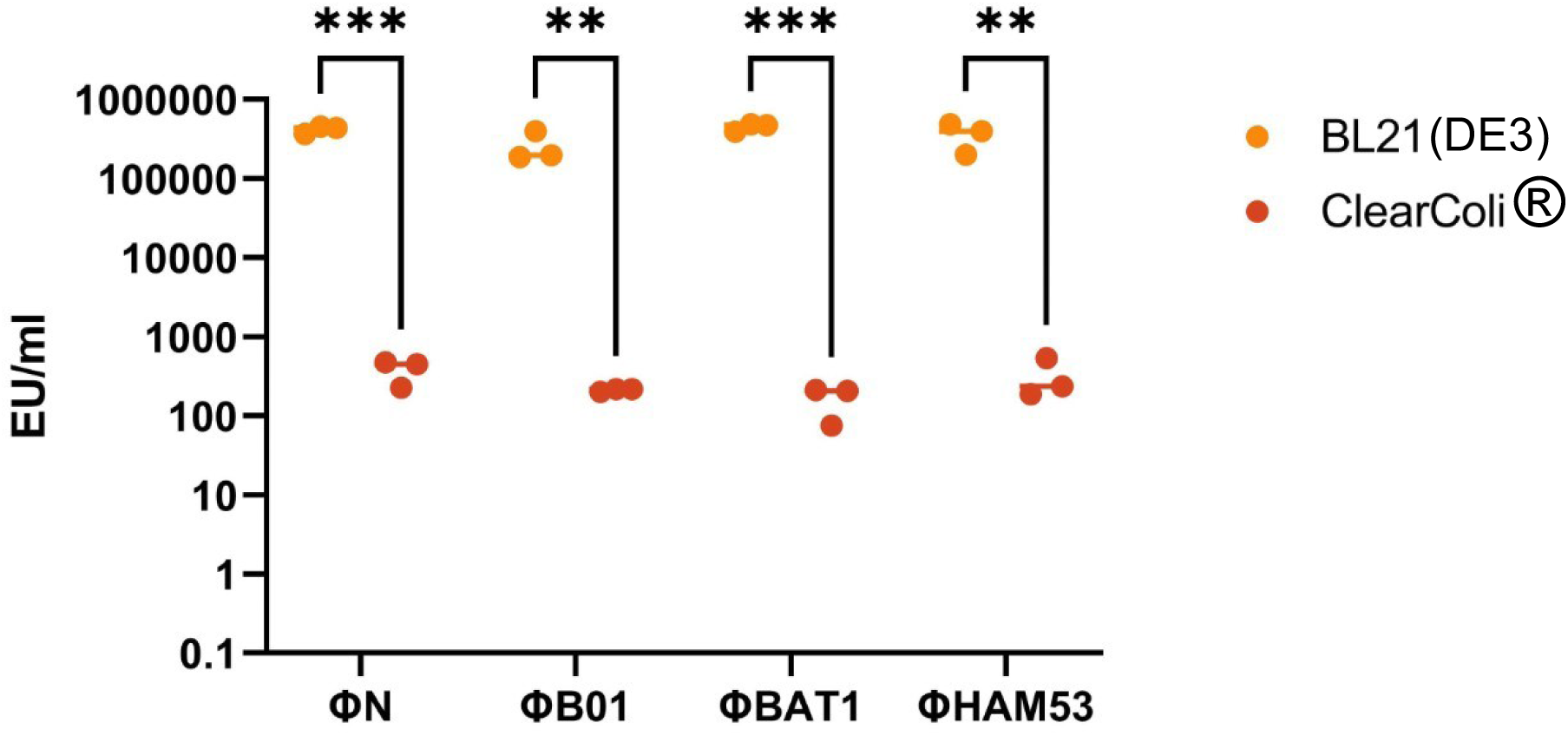
Endotoxin content of ClearColi® produced phage lysates compared to BL21(DE3). Second passage phage lysates were analysed in biological triplicate. Mean BL21(DE3) and ClearColi® lysate endotoxin levels were compared with a multiple unpaired t-test, all values were found to have significance difference, at least *P* < 0.05.

### Genetic expansion of ClearColi® phage susceptibility through receptor addition

Interestingly, LUC4 activity was observed in the parent lineage BL21(DE3) strain, despite the reported absence of its primary receptor, OmpC, in *E. coli* B-strains (26). This likely stems from the inherent genomic plasticity of BL21(DE3); *ompC* in B-strains is frequently inactivated by an upstream IS1 insertion sequence, but this leaves the possibility of an excision and recombination event restoring a functional protein (35).

Notably, this ‘leaky’ LUC4 activity phenotype was never observed in ClearColi® (**Fig. 5a**), necessitating the stable genomic integration of OmpC for LUC4 phage production. The *ompC* region including promoter *micF*, was cloned from EC958, the original isolation strain for LUC4, and allelic exchange used to replace this region in ClearColi®. On addition of the EC958 *ompC*, LUC4 is able to infect ClearColi® where it previously could not, at similar levels to the wildtype control *ompC* in EC958 (**Fig 5a**). In addition, the only other phage that lost activity between BL21(DE3) and ClearColi® (**Fig. 2**, **Table S1**), AB19 and GEO, also gained activity on the addition of the OmpC receptor (**Figs 5b and 5c**).

**Fig. 5.**
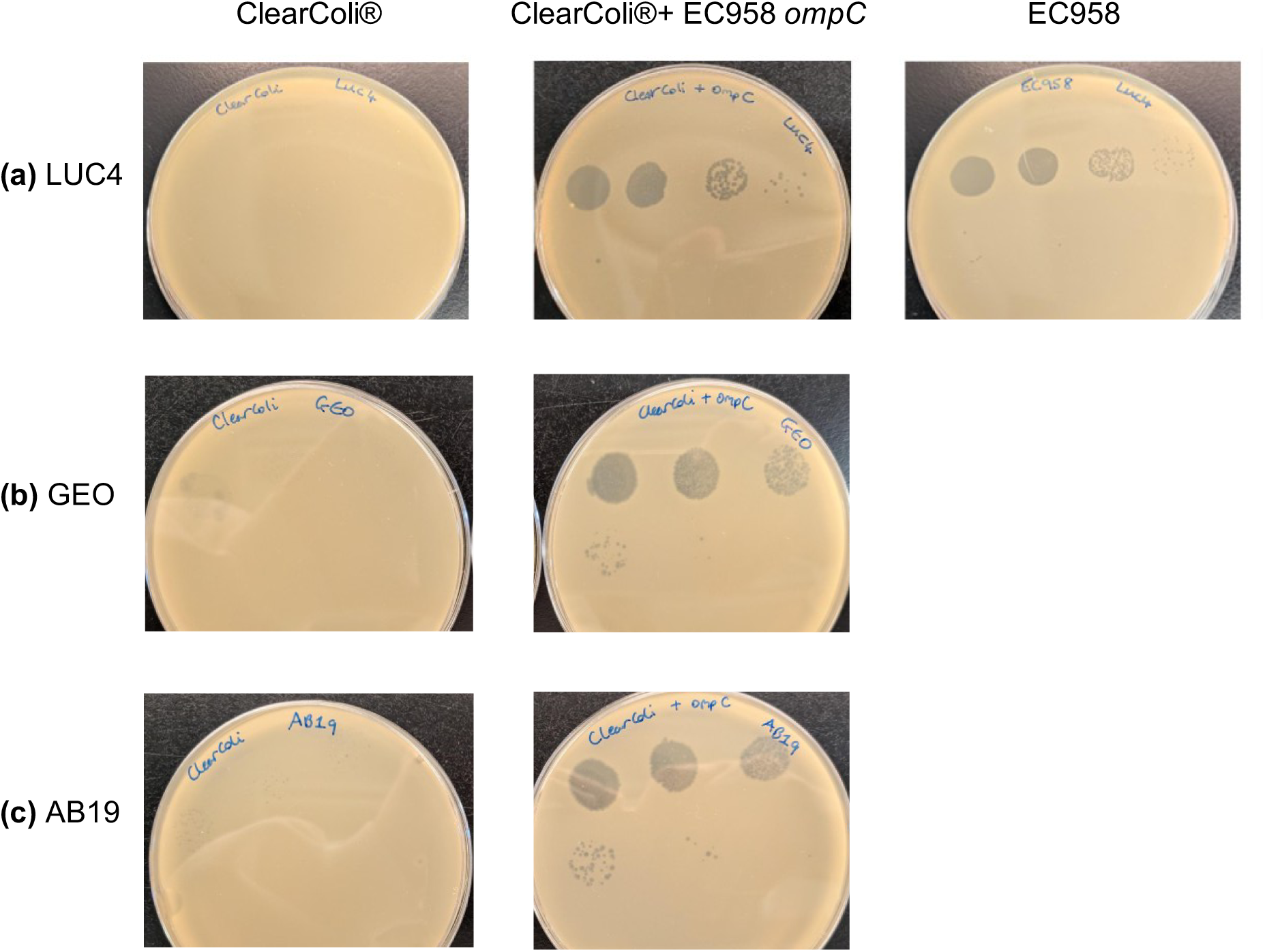
Addition of EC958 *ompC* to ClearColi® to expand phage production capacity. **a**) LUC4 10-fold serial dilution showing no successful phage infection on ClearColi® but clear evidence of plaques on ClearColi® + EC958 *ompC* and EC958 control. **b**) Phage GEO 10-fold serial dilution showing limited phage infection on ClearColi® but clear evidence of plaques on ClearColi® + EC958 *ompC.* **c**) Phage AB19 10-fold serial dilution showing limited phage infection on ClearColi® but clear evidence of plaques on ClearColi® + EC958 *ompC*.

### ClearColi® engineered to express OmpC was utilised to produce LUC4 phage for an *in vivo* safety trial

To assess the innate immune response to potential endotoxin contaminants, we monitored key biomarkers including IL-1, IL-6, IL-8 and TNF-α after administration of LUC4 phage prepared in ClearColi® + EC958 *ompC*, to three healthy pigs for an *in vivo* safety trial (**Fig. S2a**). Following intravesical phage delivery, the pigs were monitored continuously during anaesthesia recovery and for the first two hours post-procedure; no deviations in respiratory rate, behaviour, or sustenance consumption were noted. Longer-term monitoring, including body temperature (**Fig. S2b**) over 48 hours revealed no clinical abnormalities. Furthermore, serum levels of TNF-α, IFN-γ, IL-4, IL-6, IL-10 and IL-1β remained below the limits of detection, while IFN-α, IL-12/IL23p40, IL-8 concentrations showed no increase from the baseline (T_0_) (**Fig. 6**). Levels of IFN-α in pig 1 had a reduction after administration of the phage therapeutic. These preliminary results indicate that transurethral instillation of LUC4 produced in the engineered ClearColi® strain is well-tolerated in a porcine model, supporting the potential of this platform for the safe production of clinical-grade phage therapeutics.

**Fig. 6.**
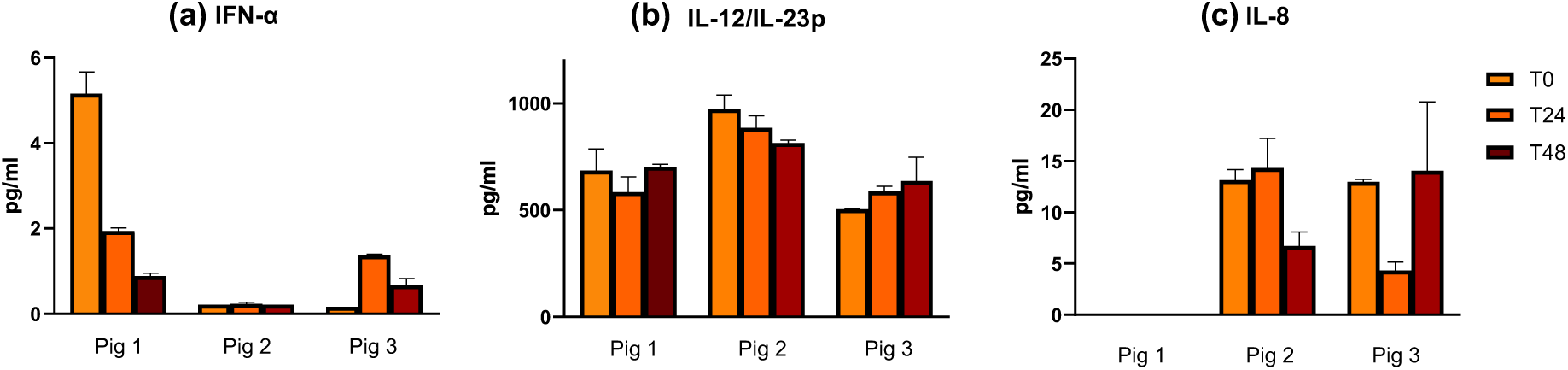
Cytokine levels at three timepoints after intravesical administration of LUC4 phage therapeutic during the *in vivo* safety trial. **a**) Levels of IFN-α in pig 1 reduced after administration of phage therapeutic. Pigs 2 and 3 had no changes in cytokine level. **b**) No large increases from T_0_ were seen in IL-12/IL-23p40 in any of the three pigs. **c**) No large increases from T_0_ were detected in IL-8 in pigs 2 and 3. Pig 1 levels were below the detection limit.

## Discussion

The escalating threat of antibiotic resistance, has revitalised commercial and academic interest in bacteriophage-based therapies. However, a primary safety hurdle is the potential for a systemic inflammatory response triggered by microbial-associated molecular patterns, in particular the Lipid A component of lipopolysaccharide (LPS), or endotoxin, present in phage preparations. This study demonstrates the utility of a detoxified *E. coli* platform for the production of safe, low-endotoxin phage lysates. While ClearColi®, was originally engineered for recombinant protein expression, we provide a proof-of-principle for its transition into the phage therapy pipeline.

Consistent with previous reports (19) we confirmed that ClearColi® exhibits a growth defect compared to the BL21(DE3) parent lineage. While this necessitated extended incubation times to reach equivalent optical densities for comparative assays, the increased doubling time had negligible impact on final phage titres. Although, parameters such as Efficiency of Plating and burst size exhibited phage dependent variability, neither host strain was consistently superior in terms of viral output. Consequently, the primary differentiator between these two strains lies in the endotoxin content of the resulting lysates rather than raw productivity. While commercial scaling would require further process optimisation for specific phage (36), our results establish ClearColi® as a high-yielding, biologically safe alternative to traditional production hosts.

While ClearColi® effectively minimises endotoxin levels, its reduced LPS profile presents a challenge for phage production, as many phage utilise LPS, often in conjunction with surface proteins such as OmpC, for cell recognition and entry (37–39). Although BL21(DE3) parent lineage has truncated LPS lacking the O-antigen and containing fewer core heptoses and hexoses than other *E. coli* strains (40,41), it probably remains more complex than the LPS of ClearColi^®^ which has been further modified by the removal of the acyl chains in the Lipid A moiety. However, only three phage had reduced infection capacity in ClearColi®, compared to BL21(DE3). For phage LUC4 this was explained with the knowledge that the receptor of LUC4 is OmpC which is differentially expressed in our parental lineage BL21(DE3) strain and ClearColi®. The other two phage that lost activity, GEO and AB19 had not had their receptors defined but activity of these phage was also observed with the addition of OmpC to ClearColi® providing strong evidence they also utilise OmpC as a receptor. Our analysis identifies LPS as a primary determinant for a subset of phage, where resistance is uniform across both strains (D11AU, F1, ALD2, GWF). The SNPs associated with resistance in *waaG* and *rfaC/F* clusters underscores a requirement for conserved outer LPS core for these phage. Two highly related broad range phage Nea2 and Chap1 do not have activity in BL21(DE3). Based on DNA sequence analysis and established taxonomic markers, phage Nea2 and Chap1 are classified as members of the genus *Tequatrovirus* (family Straboviridae); while our mutational analysis identified OmpA as the primary receptor, members of this genus typically exhibit a dual-receptor requirement, necessitating adsorption to both LPS and an outer membrane protein (42) which adds them to the subset requiring intact LPS for infection. Interestingly, Chap1 regains activity in ClearColi^®^ suggesting the severe truncation to Lipid IVa may remove the need for the multistep adsorption process and allows direct interaction with the now highly exposed OmpA. However, the overall loss of a subset of LPS-dependent phage highlights that to maximise the utility of a production host for diverse therapeutic pipelines, a strain that maintains the full structural complexity of the LPS core while strictly minimising Lipid A endotoxicity will be required to ensure propagation of a broad spectrum of LPS-dependent and protein targeting phage.

The implementation of endotoxin-reduced *E. coli* strains for the production of safe phage therapeutics necessitates a versatile host platform capable of producing the diverse lytic phage required for effective clinical cocktails. Our successful expansion of the ClearColi^®^ host range through the heterologous expression of the UPEC strain EC958 *ompC* demonstrates the potential for bespoke phage production. While the discrepancy in *ompC* expression between our BL21(DE3) lineage and the ClearColi^®^ background initially limited LUC4, GEO and AB19 production, the restoration of this receptor proved that ClearColi^®^ can be engineered to support porin-dependent phage without compromising its specialised membrane characteristics.

This modular approach allows for the targeted addition of specific receptors, such as those from clinical isolates like UPEC, to create production hosts tailored to the most therapeutically relevant phage. By decoupling the receptor requirement from the background endotoxin profile, we can generate high-titre lysates that meet stringent safety standards with minimal downstream processing. The subsequent porcine safety trial of these LUC4 lysates further validates that the genetic manipulation required to expand the host range does not interfere with the primary utility of ClearColi^®^: the provision of a safe, low endotoxin product suitable for therapeutic applications. Ultimately this strategy suggests that ClearColi^®^ could be further modified with a suite of common phage receptors (e.g., BtuB) creating a versatile library of endotoxin reduced hosts ready to produce ‘off-the-shelf’ cocktails for treating infections like UTIs.

Beyond receptor availability, the genomic landscape of the host strain must be strictly characterised to ensure the safety and efficiency of the production process. The presence of endogenous prophage or antimicrobial resistance genes within the host genome poses a risk of horizontal gene transfer; specifically, the packaging and transduction of virulence factors or resistance markers into the therapeutic lysate which could inadvertently exacerbate the infection in the patient. Furthermore, the presence of innate phage defence mechanisms, such as CRISPR-Cas or Restriction-Modification systems, can severely restrict the breadth of phage a single strain is capable of propagating. Therefore, future development of a wider range of detoxified production strains should prioritise a ‘reduced-genome’ approach and ensure detrimental genetic elements are either naturally absent or are removed to produce a genetically inert environment for safe therapeutic manufacturing.

As the UK formalises guidelines for phage therapy (11), production standards are expected to align with international limits of 5 EU/kg/hr (43,44), necessitating precision in endotoxin quantification. While the limulus amebocyte lysate (LAL) assay remains the historic gold standard (45–47), there is a critical gap in its utility for modified strains. Alternative compendial tests, such as recombinant Factor C (rFC) fluorescence assay (Pyrosmart Nextgen®), offer improved sensitivity and sustainability (48) with no activation of unspecific beta glucans, which cause interference in traditional LAL assays (49) yet they otherwise function identically to the LAL enzymatic cascade. Crucially, both LAL and rFC assays primarily detect the structural presence of the Lipid A moiety and potentially won’t detect the differing acylation patterns of a modified strain that dictate human immune activation.

In contrast, our use of the SEAP-based TLR4 reporter assay (EndotoxDetect™, InvivoGen) provides a direct measure of biological pyrogenicity (50). This method confirms that while Lipid IVa may still be chemically detectable by traditional assays, it fails to trigger the TLR4-mediated inflammatory response characteristic of standard LPS. This discrepancy presents a regulatory challenge: industrial scale-up remains tethered to LAL-based metrics, even when those metrics do not reflect the true safety profile of a modified lysate. However, the significant reduction in biological activity demonstrated in ClearColi^®^-produced lysates offers an economic advantage. By utilising a ‘detoxified-by-design’ production host, the requirement for costly and yield-limiting downstream purification steps, such as octanol extraction or ion exchange chromatography, is drastically reduced. For industrial scale phage manufacturing, the integration of strains like ClearColi^®^could fundamentally lower the overhead of compliance, providing a more efficient pathway for producing high titre, safe phage therapeutics for complex infections.

While *in vitro* assays provide essential data, the safety of phage therapeutics must ultimately be demonstrated within a complex biological system. To this end, our porcine safety trial served as *in vivo* validation confirming that the low-endotoxin lysates produced from ClearColi^®^ did not trigger an adverse systemic immune response. Pigs are an ideal model for this assessment due to their physiological and immunological similarities to humans, particularly regarding their sensitivity to cytokine storms. It is important to note that this trial utilised intravesical administration, reflecting the intended clinical route for treatment of UTIs. Unlike intravenous (IV) administration, which requires strict adherence to systemic endotoxin limits, the intravesical route allows for the assessment of safety in the context of the uroethelial barrier. The absence of fluctuations in pro-inflammatory cytokines following administration indicates within a mucosal environment, ClearColi® lysates are systemically inert. This mirrors the results of previous studies that have demonstrated the safety of purified phage lysates in both animal and human clinical cases (51). This study demonstrates using a genetically detoxified strain is sufficient to achieve safety for localized administration without the need for aggressive chemical de-pyrogenation.

This study establishes a clear proof of principle that endotoxin-reduced *E. coli* strains can be used for the streamlined production of ‘safe’ phage therapeutics without significantly compromising viral titres or host range. While ClearColi® is a commercial platform requiring licensing for product development, its utility demonstrates the potential of genetically detoxified hosts. The structural limitations observed, specifically the lack of a complete lipopolysaccharide outer core and O-antigen, do restrict the propagation of some LPS-dependent phage; however, our work shows targeted receptor engineering can expand the phage host range and similar detoxification mutation strategies could be applied to different strain backgrounds.

By integrating these detoxified-by-design production strains into manufacturing pipelines the requirement for extensive downstream processing is minimised, potentially allowing for the near-immediate administration of lysates in urgent clinical situations. Ultimately, this production model offers a strategy for the future of personalised medicine, shortening the window between clinical diagnosis and therapeutic application. By utilising a curated library of endotoxin reduced production strains the transition from bench-to-bedside can be accelerated, providing a rapid, safe as well as economically scalable solution for the global challenge of antimicrobial resistance.

## Author statements

### Author contributions

Conceptualisation: A.L., D.G.; Data curation: E.W. A.P.d.l.T.; Formal analysis: E.W. A.P.d.l.T.; Funding acquisition: A.L., G.P., D.G.; Investigation: E.W., A.P.l.d.T., J.A.N., M.K., S.M.; Methodology: A.L., E.W., A.P.d.l.T., S.M., Project administration: A.L., D.G. Resources: E.W., A.P.l.d.T., J.A.N., M.K., S.M.; Supervision: D.G., A.L., G.P.; Validation: E.W.; Visualization E.W., A.P.d.l.T., A.L.; Writing - original draft: E.W., A.L., D.G.; Writing - review & editing: E.W., A.L., D.G., G.P.

### Ethical Approval

The porcine safety trial was carried out under Home Office Project License (PCD70CB48), and approved by the University of Edinburgh Animal Welfare and Ethical Review Body (AWERB).

## Acknowledgements

The authors are grateful to the staff at Easter Bush Large Animal Research and Imaging Facility (LARIF) for their invaluable assistance with the porcine safety trial. We would like to thank Professor Mark Stevens for his oversight and for conducting the animal procedures under his Home Office Project License. We also appreciate the expertise of Dr. Nicky Craig in the execution and analysis of the Luminex assays.

## Supplementary Tables and Figures

**Fig. S1.**
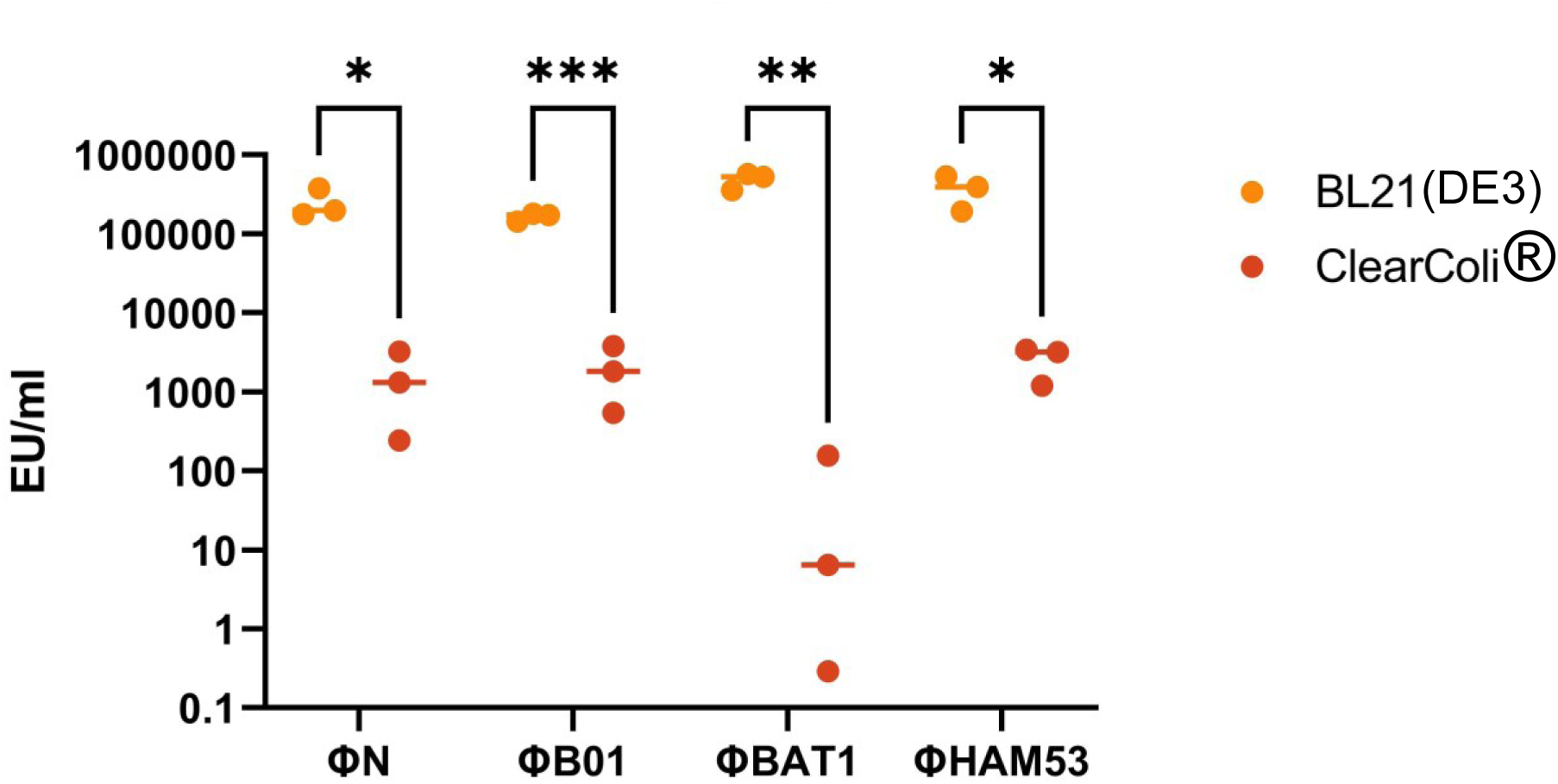
Endotoxin content of first passage phage lysates. First passage phage lysates were analysed in biological triplicate. Mean BL21(DE3) and ClearColi® lysate endotoxin levels were compared with a multiple unpaired t-test, B01 and BAT1 values were found to have significance difference, at least P < 0.05. Phage N and HAM53 lysates were not significantly different between BL21(DE3) and ClearColi®, likely due to carryover of endotoxin from the original host bacteriophage production strain, hence a second passage was completed (**Fig. 5**).

**Fig. S2.**
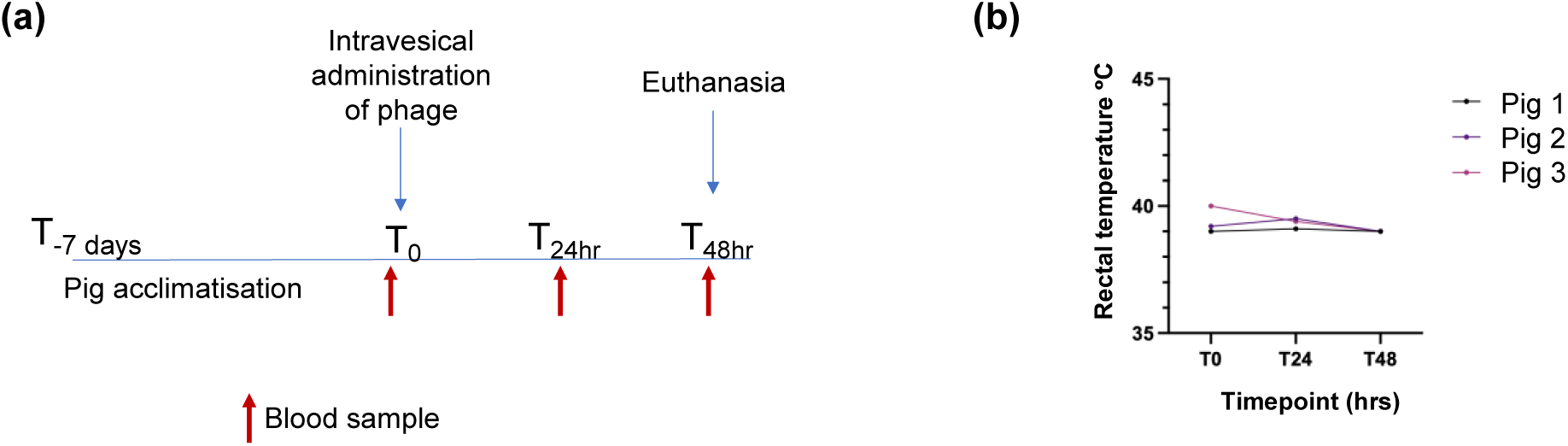
*In vivo* phage safety trial. (a) Study plan for the *in vivo* safety trial. Pigs were acclimatised in the facility for 7 days before intravesical administration of the phage preparation at time T_0_. Blood samples for cytokine monitoring were taken just prior to phage administration, at 24 hours and at 48 hours just prior to euthanasia. (b) Rectal temperatures of the pigs were obtained at each timepoint of the study.

**Table S1.** Comparison of phage activity between BL21(DE3) and ClearColi® and identified phage receptors. Area Under the Curve scores of 31 phage infecting BL21(DE3) or ClearColi®. A score of over 60 (red text and red/orange shades on the heatmap) indicates non-active phage, while lower scores of less than 60 indicate positive phage infection (yellow to green shades on the heatmap). Scores calculated using biological triplicates of 18-hours of growth data. Red text phage are classified as no activity in BL21(DE3) or ClearColi®. Blue text phage have lost activity in ClearColi® and purple text phage have gained activity in ClearColi®. Sequencing of fixed phage resistant mutants (original host strains) revealed SNPs in genes allowing prediction of phage receptors. Predicted receptors shaded in orange highlight phage that are predicted (from SNPs) to utilise LPS as a receptor. Boxes shaded grey represent receptor not yet identified.

| Phage | Phage Score |  | Predicted Receptors | Genes with SNPs |
| --- | --- | --- | --- | --- |
|  | BL21(DE3) | ClearColi® |  |  |
| NEA2 | 100 | 79 | OmpA | *ompA* |
| F1 | 100 | 86 | LPS | *rfaF* |
| D11AU | 100 | 99 | LPS | *waaG* |
| ALD2 | 98 | 97 | LPS | *waaG, rfaC* |
| OIL | 97 | 92 | BtuB | *btuB* |
| TB69 | 90 | 91 |  |  |
| D3 | 87 | 89 | Capsule | *kpsC* |
| GWF | 86 | 81 | LPS | *rfaY* |
| CHAP1 | 85 | 44 | OmpA | *ompA* |
| D4 | 51 | 53 | LPS | *hdiE* |
| UFA5s | 50 | 42 |  |  |
| E4 | 49 | 35 | LPS and OmpA | *rfaH, wzxC, rfaF, ompA* |
| B01 | 46 | 46 | LPS | *rfaC* |
| GEO | 42 | 64 | OmpC* |  |
| LUC4 | 41 | 89 | OmpC | *ompC* |
| RV2 | 36 | 27 | LPS | *rfaC* |
| A01 | 36 | 7 | FhuA | *fhuA* |
| ALD1 | 31 | 7 | FhuA | *fhuA* |
| ALDI98 | 31 | 16 | FhuA | *fhuA* |
| HAM3 | 29 | 16 |  |  |
| A | 28 | 15 | OmpX, LpS | *ompX, rfaG* |
| AB19 | 28 | 74 | OmpC* |  |
| P | 24 | 19 | OmpX | *ompX* |
| LEG | 23 | 11 | LPS | *ggt, pgm* |
| N | 21 | 10 | OmpX, LPS | *ompX, rfaG* |
| E | 21 | 10 | OmpX, LPS | *ompX, hdiE* |
| BAT1 | 18 | 10 | Tsx | *tsx* |
| TB54 | 18 | 8 |  |  |
| HAM53 | 17 | 8 | OmpA | *ompA* |
| BOW | 14 | 10 | LPS | *rfaD, wzzB* |
| FAT | 11 | 8 | LPS | *rfaP* |

\*receptor determined by activity on ClearColi® +ECompC but no activity on ClearColi® (Fig. 5)

